# Temporal Mechanisms of T-Cell Fate Decisions under Immune Checkpoint Blockade Resolved by CanonicalTockySeq

**DOI:** 10.64898/2026.03.10.710825

**Authors:** Jehanne Hassan, Omnia Reda, Nobuko Irie, Malin Pedersen, Shane Foo, Lizzie Appleton, Il-mi Okazaki, Taku Okazaki, Yorifumi Satou, Kevin Harrington, Alan Melcher, Masahiro Ono

## Abstract

In cancer immunotherapy, T-cell functional states change dynamically, and no single marker or gene fully defines them. Static molecular profiles may therefore be insufficient to resolve these states. Capturing signalling history together with ongoing activity could provide a temporal framework for their dissection. Here, we developed CanonicalTockySeq, which integrates a molecular clock of T-cell receptor (TCR) signalling based on the Fluorescent Timer reporter Nr4a3-Tocky (Timer-of-cell-kinetics-and-activity) with scRNA-seq to establish an experimentally anchored temporal reference. Using landmark Tocky fractions as biological ground truths, CanonicalTockySeq constructs a transcriptomic manifold in canonical space with conical geometry. In a murine melanoma model treated with combination anti-PD-L1 and anti-CTLA-4, this framework separates temporal progression (geodesic angle) from signalling strength (radial intensity), enabling time-resolved analysis of gene-expression programmes at single-cell resolution. Application to scRNA-seq data from patients with melanoma further identifies distinct temporal programmes associated with clinical response. These analyses indicate that effective combination immunotherapy is associated with decreased persistence of antigen engagement, suppression of exhaustion-associated TCR signalling programmes, and maintenance of progenitor-like features linked to durable antitumour responses. Collectively, our findings identify temporal-state control as a key component of immunotherapy outcome and establish CanonicalTockySeq as a framework for resolving T-cell response states in vivo.

## Introduction

Single-cell RNA sequencing (scRNA-seq) has revolutionized our understanding of cellular heterogeneity, yet it remains fundamentally constrained by its nature as a destructive, cross-sectional “snapshot” ^1, 2, 3^. In the context of tumour immunology, these snapshots provide a high-dimensional view of T-cell states ^4^ but fail to capture the critical temporal history of T-cell receptor (TCR) engagement ^5^. TCR signalling, indeed, has dominant impacts on T-cells’ fates ^6, 7^, including whether a T cell becomes a potent effector or enters a state of terminal exhaustion ^8, 9, 10, 11^.

While computational frameworks like pseudotime trajectory analysis ^12^ and RNA velocity ^13^ attempt to infer cellular “progress”, they are inherently limited ^3^. Pseudotime identifies potential lineages by ordering cells based on transcriptomic similarity, yet it lacks an absolute temporal anchor, making it prone to circular reasoning in chaotic tumour environments ^14^. RNA velocity provides a predictive vector of cell states by modelling the kinetics of mRNA splicing ^13, 15^; however, its assumption of constant transcription and degradation rates is often invalidated by the rapid metabolic shifts of T-cell activation ^16, 17^.

Furthermore, while physical time-stamping and nascent RNA methods, such as scGRO-seq ^18^ and metabolic pulse-chase labeling (e.g., scSLAM-seq) ^19^, can elucidate acute, time-dependent changes in transcription, they often require controlled ex vivo conditions or invasive labelling that is difficult to scale in vivo. Zman-seq introduces fluorescent “*time stamps*” into circulating immune cells to track their infiltration history ^20^, but is not suitable for analysing a continuous, endogenous record of signalling events. Consequently, a robust methodology for capturing continuous, endogenous temporal resolution anchored to TCR signalling history and timing has yet to be established.

To bridge this gap, the Timer-of-cell-kinetics-and-activity (Tocky) system offers a unique approach by utilizing a fluorescent Timer protein ^6^, which matures from an unstable blue-fluorescent form to a stable red-fluorescent form with predictable kinetics ^21, 22^. Unlike purely computational pseudotime or metabolic labelling methods, the Nr4a3-Tocky system provides a continuous, high-resolution molecular chronometer anchored to the “ground truth” of T-cell receptor (TCR) signalling. Because Nr4a3 transcription is negligible in resting T cells and is strictly induced upon TCR engagement ^6, 23^, the blue-to-red maturation process enables the physical identification and isolation of cells at precise stages of their signalling history. Previously, we established a canonical analysis framework for flow cytometric data using Canonical Correspondence Analysis (TockyCCA) ^24^. However, methods to elucidate time-dependent gene expression profiles with an empirical temporal resolution using Tocky are yet to be established.

The application of the Nr4a3-Tocky system to tumour-infiltrating lymphocytes (TILs) addresses two key gaps in immuno-oncology. First, it separates differentiation state from TCR signalling dynamics, enabling tests of whether exhaustion represents a distinct differentiation trajectory ^25^ or a consequence of cumulative signalling over time ^9^. Second, it provides a high-resolution temporal readout to analyse how Immune Checkpoint Blockade (ICB) reshapes baseline anti-tumour T-cell responses ^26, 27^. While ICBs targeting the PD-1/PD-L1 and CTLA-4 axes have fundamentally shifted the treatment landscape by reinvigorating tumour-reactive, but dysfunctional, T cells ^28^, the temporal mechanisms by which initially reactive T cells become exhausted, and how this process is reversed over time, remain unclear ^29, 30^. Current evidence suggests that ICB primarily acts on a subset of “progenitor exhausted” T cells ^31, 32^, yet it remains unclear whether these cells are caught in a specific signalling window or if therapy induces a genuine temporal “reprogramming” of the T-cell life cycle ^33^. Defining the time domain of the T-cell response is, therefore, essential for optimizing therapeutic timing and identifying the precise molecular checkpoints that govern successful reinvigoration versus terminal failure.

Building on our series of Tocky-based methodological studies, we developed CanonicalTockySeq, a multidimensional extension of Tocky theory that transforms the Timer’s temporal logic into a transcriptomic manifold using experimentally defined Tocky signatures. Using this innovative method, we analysed how immunotherapies modify the temporal programme of anti-tumour T cells.

## Results

### Establishment of Experimentally Anchored Tocky-Time Trajectory Analysis for Single-Cell Transcriptomics

To rigorously quantify the temporal dynamics of cellular differentiation and activation in vivo, we utilised the Nr4a3-Tocky system, which reports transcriptional activity through a Fluorescent Timer protein that spontaneously matures from an unstable blue-fluorescent form ( *t*1/2 ≈ 4 h ^22^) to a stable red-fluorescent form (*t*_1/2_ ≈ 122 h ^22^). Flow cytometric analysis resolves these dynamics in a two-dimensional “Timer Space” (Blue vs. Red fluorescence), where the position of each cell reflects its transcriptional history (**Figure 1a**). By converting these Cartesian coordinates into polar coordinates, we defined “Tocky Time” as the angle (θ) relative to the Red axis ^34^. This metric stratifies cells into five continuous signalling stages: New (recently initiated signalling), NPt (New-to-Persistent transitioning), Persistent (sustained signalling), PAt (Persistent-to-Arrested transitioning), and Arrested (signalling ceased) ^6, 35^. Functionally, this trajectory resolves two domains: a “Time Sequence” domain (New to Persistent), governed by Timer maturation following transcriptional burst, and a “Frequency” domain (Persistent to Arrested), where sustained or repetitive signalling drives accumulation of the stable Red form ^6^.

**Figure 1.**
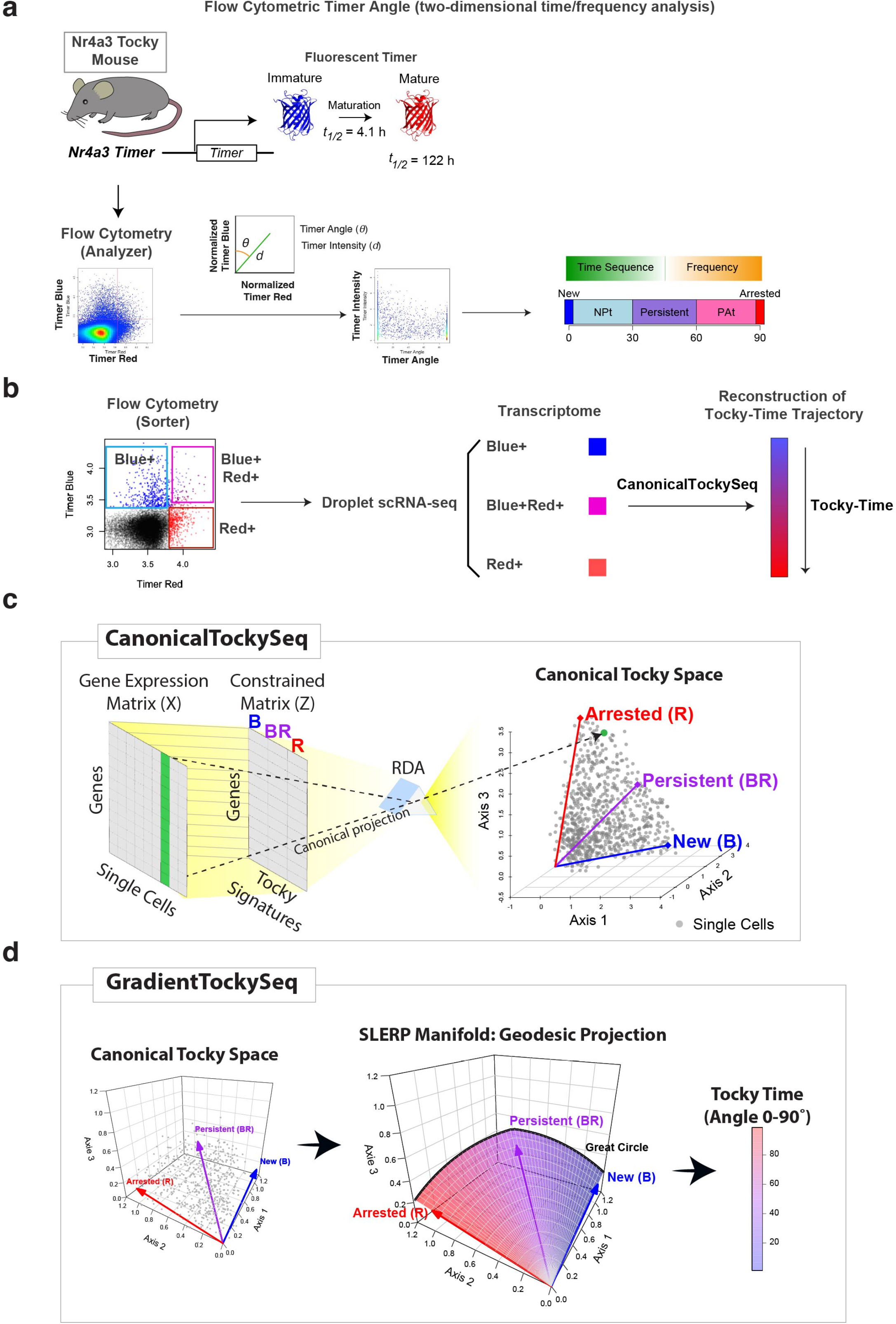
Establishment of the CanonicalTockySeq Framework for Multidimensional Analysis of Temporal Differentiation. (a) The Biological Basis of The Nr4a3-Tocky Timer. Schematic illustrating the Nr4a3-Tocky system, where transcriptional initiation triggers the production of a Fluorescent Timer protein that matures from an unstable Blue form (*t*_1/2_ ≈ 4 h) to a stable Red form (*t*_1/2_ ≈122 h). Flow cytometric analysis resolves these dynamics in a two-dimensional “Timer Space”. The original Tocky Angle is defined trigonometrically relative to the Red axis, stratifying cells into developmental stages from “New” (recent initiation) to “Arrested” (cessation of signalling). (b) Experimental workflow for establishing biological ground truths for transcriptomic analysis. Nr4a3-Tocky T cells stimulated in vivo are physically isolated via Fluorescence-Activated Cell Sorting (FACS) into three discrete reference populations based on their Timer locus positions: Blue+ (B), Blue+Red+ (BR), and Red+ (R). These populations are subjected to droplet-based single-cell RNA sequencing (scRNA-seq) to define Tocky-state gene signatures, with the aim of reconstructing continuous Tocky-Time trajectory utilising the CanonicalTockySeq algorithm. (c) Schematic of the CanonicalTockySeq methodology. A supervised Canonical Redundancy Analysis (RDA) is employed, where the single-cell gene expression matrix (*x*) is constrained by an explanatory matrix (***z***) derived from the reference Tocky signatures (B, BR, R). Through Canonical projection, high-dimensional transcriptomic data is mapped onto a low-dimensional manifold strictly governed by the Timer phenotypes. The resulting 3D “Canonical Tocky Space” visualizes individual cells (dots) preserving their biological heterogeneity, oriented by biplot vectors representing the Blue, Blue-Red, and Red reference Tocky signatures. (d) Schematic of GradientTockySeq for trajectory reconstruction and quantification. A continuous, piecewise geodesic trajectory is constructed by connecting the landmark biplot vectors—New (B) → Persistent (BR) → Arrested (R), using Spherical Linear Interpolation (SLERP) ^38^. Individual cells are projected onto this manifold to derive two metrics: *Tocky Time*, the angular position (θ) along the geodesic path normalised from 0° to 90°; and *Tocky Intensity*, the normalised radial magnitude representing the collective strength of the Tocky-associated transcriptional program.

To extend this temporal resolution from a single reporter protein to the entire transcriptome, we developed the `CanonicalTockySeq` framework (**Figure 1b**). We physically isolated Nr4a3-Tocky T cells into three discrete populations using fluorescence-activated cell sorting (FACS), including Blue^+^Red^-^ (New), Blue^+^Red^+^ (Persistent), and Blue^-^ Red^+^ (Arrested), and subjected them to droplet-based single-cell RNA sequencing (scRNA-seq). These sorted populations served as biological ground truths, providing the transcriptomic constraints necessary to reconstruct a continuous “Tocky Time” trajectory solely from gene expression data.

To integrate the temporal dynamics of cellular differentiation with high-dimensional transcriptomic states, we developed CanonicalTockySeq, a multidimensional extension of the Tocky Angle theory ^6, 36^. The original framework utilised Timer fluorescence maturation to define a 2D polar coordinate, successfully identifying the in vivo sequence from transcriptional initiation (New) to cessation (Arrested). Built on this work, we hypothesised that the transcriptome itself could orient this temporal metric. By incorporating the experimental analysis of reference Tocky populations (**Figure 1b**), CanonicalTockySeq transforms the Tocky Angle from a 2D fluorescence readout into a multidimensional transcriptomic manifold.

As current droplet-based scRNA-seq cannot simultaneously quantify Timer protein and transcriptomes in the same cell, we implemented a geometric reconstruction strategy that uses flow-sorted Blue (B), Blue/Red (BR), and Red (R) populations as biological landmarks to derive Tocky-state gene signatures (**Figure 1c**). CanonicalTockySeq applies the supervised canonical method, redundancy analysis (RDA) ^37^, to constrain the transcriptomic space by the Tocky signatures (New [B], Persistent [BR], Arrested [R]; **Supplementary Methods**). Individual single cells are subsequently projected onto this structure, allowing us to visualize their differentiation state while preserving biological heterogeneity (**Figure 1c**). In this representation, biplot vector direction encodes the Tocky-defined time dimension, and vector magnitude quantifies collective signature strength, reflecting TCR-signalling intensity in the context of Nr4a3-Tocky.

To convert manifold geometry into quantitative temporal metrics, we developed GradientTockySeq using Piecewise Spherical Linear Interpolation (SLERP) ^38^. The manifold is anchored by landmark vectors corresponding to the New (B), Persistent (BR), and Arrested (R) signatures (**Figure 1d, Supplementary Methods**).

While linear interpolation assumes a straight-line path between states, Piecewise SLERP generates a smooth trajectory that maintains constant angular velocity in direction space ^38^. Crucially, as the landmark vectors for B, BR, and R typically differ in Euclidean magnitude, reflecting varying transcriptional intensities, the resulting SLERP trajectory traces a piecewise logarithmic spiral. This spiral path captures the biological reality that cells do not just shift their expression profile according to time (Timer Angle) but also reflect the accumulated dynamics of total TCR signals (Timer Intensity) as they mature.

By considering rays from the origin through this path, the model provides a continuous cone-like geometric framework. Operationally, cells are mapped by maximising similarity to interpolated reference vectors along the piecewise path, yielding a temporal coordinate (Tocky Time), while vector magnitude is quantified separately as intensity. Geometrically, CanonicalTockySeq is formulated in a conical coordinate system: temporal progression is modelled on the hyperspherical angular manifold defined by landmark directions, whereas radial distance from the origin encodes signal intensity (**Figure 1d**). Complementing this angular metric, we defined Tocky Intensity by normalising the radial distance against the landmark magnitudes. This ensures that intensity reflects biologically meaningful signature strength rather than geometric scale differences inherent to the constrained space.

### In Silico Validation of CanonicalTockySeq and the Geodesic Manifold

To assess the fidelity of the GradientTockySeq method rigorously, we conducted an in silico validation using synthetic single-cell data generated along engineered trajectories. First, to identify the optimal gradient model, we first compared two geometric interpolation strategies, Quadratic Bézier interpolation and Piecewise Spherical Linear Interpolation (SLERP), under two scenarios: an ideal orthogonal trajectory and a biologically realistic skewed, non-orthogonal trajectory (**Supplementary Figure 1a–1d**). Under the orthogonal condition, both methods recovered the simulated temporal progression with high fidelity (**Supplementary Figure 1a–1b**). However, under the skewed condition, the SLERP-based geodesic path showed substantially greater robustness, maintaining near-linear agreement with the true simulated time (r = 0.97), whereas the Bézier model introduced marked sigmoidal distortion (**Supplementary Figure 1c–1d**). In parallel, SLERP provided more stable intensity normalisation across the trajectory, supporting its suitability for downstream temporal analysis in non-ideal ordination spaces (**Supplementary Figure 1d**).

We next examined whether unsupervised ordination alone could recover biologically meaningful Tocky structure. Although unsupervised PCA successfully identified the primary structure of the simulated data (**Supplementary Figure 2a**), the PCA axes themselves did not provide an intrinsic mapping to the biologically defined Tocky stages (New, Persistent, and Arrested). In contrast, CanonicalTockySeq used landmark Tocky signatures to construct a supervised canonical space in which the manifold was explicitly aligned to these biological reference states (**Supplementary Figure 2b**). This preserves the global arched structure observed by PCA while anchoring it to interpretable temporal vectors.

Using the three biologically defined poles, Blue (New), Blue-Red (Persistent), and Red (Arrested), SLERP-based gradient analysis accurately reconstructed the programmed sequence of gene activation, resolving 100 simulated genes into a coherent temporal cascade ordered by peak expression time (**Supplementary Figure 2c**). These simulations show that the CanonicalTockySeq framework can recover temporal progression with high fidelity, preserve biologically meaningful manifold geometry, and decouple differentiation timing from signal intensity before application to complex in vivo datasets.

### Elucidating the Temporal Mechanism of Antigen-Reactive CD8 T cells in Melanoma and the Impacts of Combination Immunotherapy

To validate the utility of CanonicalTockySeq in dissecting transcriptomic mechanisms in T cell responses in vivo, we analysed CD8^+^ T cells from tumour-infiltrating lymphocytes (TILs) and draining lymph nodes of Nr4a3-Tocky mice bearing B16 melanoma. Mice were treated with combination anti-PD-L1 and anti-CTLA-4 (combination) or isotype control IgGs on days 12, 14, and 16, followed by the analysis and library construction at day 18 (**Figure 2a**). This design allowed us to capture the early post-treatment transcriptional and signalling-state response of T cells under combination immunotherapy. Flow-sorted Tocky landmark fractions, New (Blue^+^Red^−^), Persistent (Blue^+^Red^+^), and Arrested (Blue^−^Red^+^), were used to perform droplet-based scRNA-seq. The data were further subjected to in silico sorting to purify CD4^+^ and CD8^+^ T cells, to which the CanonicalTockySeq pipeline was applied using the data from the three Tocky landmarks as explanatory vectors to project cells into a three-dimensional Canonical space.

**Figure 2:**
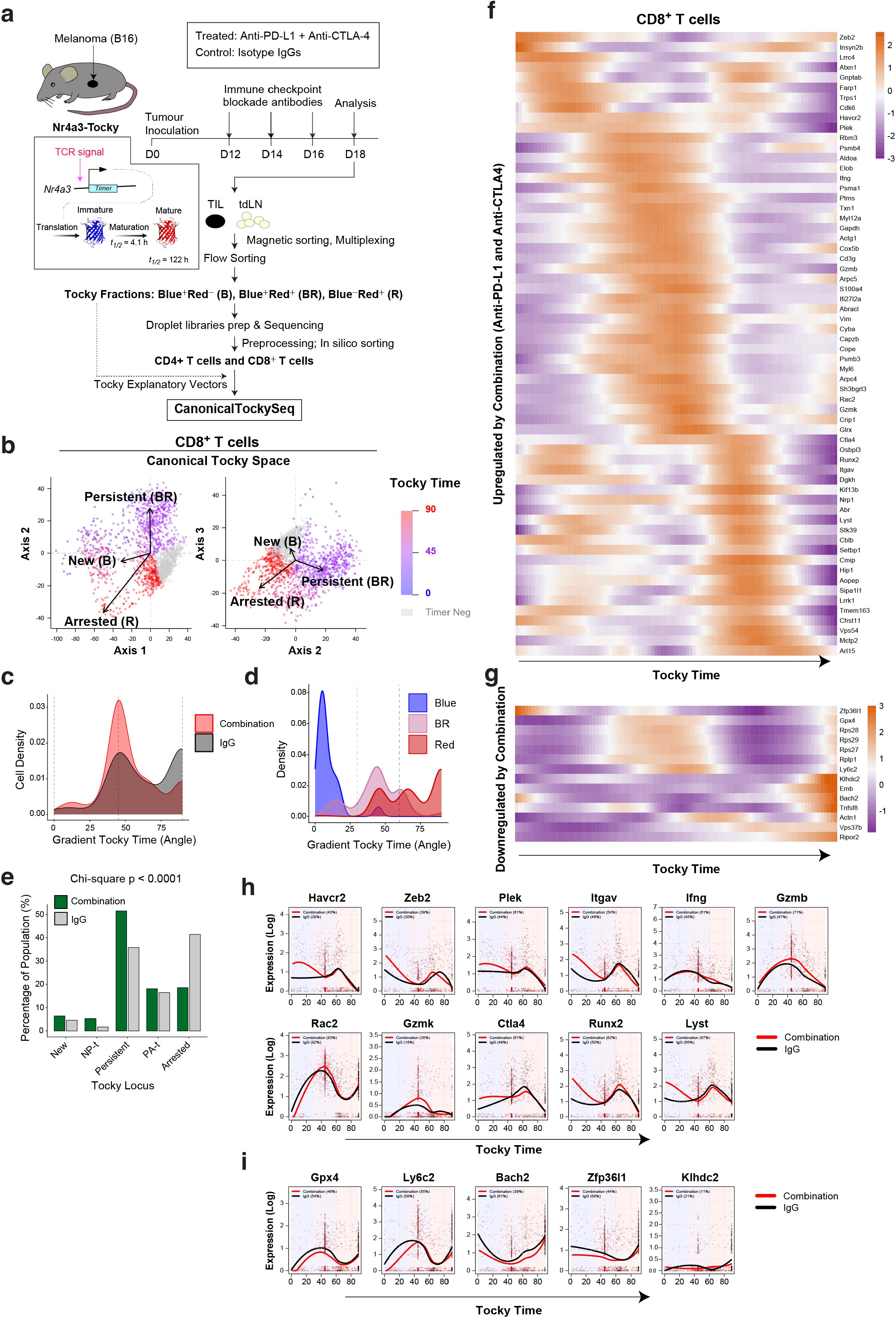
CanonicalTockySeq reveals the temporal dynamics of checkpoint blockade response in CD8^+^ T cells. (a) Experimental design: B16-F10 melanoma-bearing Nr4a3-Tocky mice were treated with anti-PD-L1 and anti-CTLA-4 (Combination) or Isotype IgG. TILs were harvested on day 16. (b) Canonical Tocky Space analysis of CD8+ TILs, showing the progression from New (Blue) to Arrested (Red) phenotypes along the Timer trajectory. (c) Angular density plot showing the massive accumulation of combination-treated cells (Red fill) in the intermediate trajectory ( ≈ 45°) compared to IgG controls (Grey fill). (d) Validation of manifold fidelity: Distribution of calculated Gradient Tocky Times for experimentally sorted Blue, BR, and Red populations. (e) Tocky Locus quantification confirming the significant expansion of the Persistent fraction in the treatment group (Green bars) versus IgG (Grey bars). (f-g) Temporal heatmaps of genes upregulated (f) or downregulated (g) by combination therapy, ordered by peak expression timing. (h-i) Kinetic expression profiles of key effector (h) and stemness (i) regulators. Combination therapy (Red lines) drives earlier/higher induction of *Gzmb* and *Ifng* while suppressing the memory guardians Bach2 and Zfp36l1 compared to IgG (Black lines).

Using the sorted CD8^+^ T cell data, CanonicalTockySeq applied RDA to project the full scRNA-seq dataset onto the first three canonical axes, thereby constructing a continuous differentiation manifold (**Figure 2b**). The Tocky explanatory vectors, i.e. New (B), Persistent (BR), and Arrested (R), define the trajectory’s poles, with individual cells distributed along this path according to their Gradient Tocky Time (angle). In the Canonical Tocky model constructed for CD8^+^ T cells, Tocky-Time distribution analysis shows that cells with a “New” phenotype (blue, angle ≈ 0°) cluster around the ‘New (B)’ vector, while those with a “Persistent” phenotype (purple, angle ≈ 45°) align with the ‘Persistent (BR)’ vector (**Figure 2c**). The trajectory terminates with cells in the “Arrested” state (red, angle ≈ 90°), which are concentrated at the pole of the ‘Arrested (R)’ vector.

To confirm that the CanonicalTockySeq projection accurately captured the biological maturation of the Nr4a3-Tocky signal, we cross-validated the calculated Gradient Tocky Time (Angle) against the original sorted fractions. The trajectory landmarks showed high fidelity to the experimental sorting labels (**Figure 2d**). Cells sorted as “Blue” (new antigen experience) were tightly confined to the early trajectory (Angles 0–25◦). The “BR” fraction occupied the intermediate zone, while the “Red” fraction exhibited a bimodal distribution: a subset of cells overlapped with the BR peak at ∼ 45◦, identifying a “late-persistent” state, while the majority accumulated at the terminal end of the trajectory (>75◦). This confirms that the manifold successfully resolves the continuous nature of differentiation that is often obscured by discrete flow cytometry gates.

We next used this validated trajectory to quantify the effect of combination immunotherapy (anti-PD-L1 plus anti-CTLA-4), which was selected for scRNA-seq as the principal treatment condition to enable deep mechanistic profiling without substantially increasing the number of experimental groups and sequencing libraries, using the Tocky Locus approach (**Figure 2e**). A chi-square test confirmed a non-random redistribution of CD8^+^ T cells across the five Tocky Loci (p < 0.0001). In IgG-treated controls, CD8^+^ TILs predominantly progressed to the terminal Arrested phase, whereas in the combination-treated group the Persistent phase expanded significantly, accounting for ∼50% of CD8^+^ T cells. Given that this analysis was performed 2 days after the final treatment dose, these findings suggest that, at an early post-treatment time point, a large fraction of tumour-reactive CD8^+^ T cells remain actively engaged with target antigens under combination immunotherapy.

To deconvolve the temporal architecture of the therapeutic response, we utilised a Tocky-Time heatmap approach, which aligns genes sequentially according to their peak expression timing along the trajectory. This method visualises gene expression dynamics as a continuous activation or silencing cascade, allowing us to resolve the precise timing of transcriptional transitions within the validated manifold (**Figure 2f–g**). This approach orders cells by their Gradient Tocky Time, revealing the precise sequential timing of gene activation and silencing as experimentally measured by Tocky and reconstructed by the CanonicalTockySeq manifold.

**Figure 2f** displays the “activation cascade” of genes significantly upregulated by combination therapy. Sorting these genes by their peak expression time revealed a strictly ordered differentiation program. The cascade initiated with immediate-early genes associated with effector fate commitment (e.g., *Zeb2, Atxn1*) and early-onset immune regulation (e.g., *Havcr2*), which were induced at the onset of the trajectory (New-to-Persistent transition). These were accompanied by drivers of structural reorganization and synapse priming (e.g., *Farp1*), indicating that the transcriptional program for both functional identity and physical motility is established prior to the acquisition of full cytotoxic capacity. Following this, the core cytotoxic machinery, including *Gzmb, Ifng*, and *Gzmk*, reached peak intensity in the intermediate Persistent phase (Angle ∼ 45◦), coincident with the population density accumulation.

Conversely, **Figure 2g** visualises the “silencing cascade” of genes downregulated by the therapy. This panel highlights the progressive loss of the stemness programme. Notably, the combination therapy repressed key memory regulators such as *Bach2* and *Zfp36l1*, which were active in the “New” and “Arrested” phases but silenced as cells entered the Persistent phase.

Trajectory-resolved analysis of genes induced by combination therapy (**Figure 2h**) further refined this observation, showing that combination therapy not only increased effector-gene expression but also altered its temporal pattern along the trajectory. Combination-treated cells expressed *Gzmb* and *Ifng* at significantly higher levels and with an earlier peak timing (shifted to the left) compared to IgG controls. This accelerated effector burst was accompanied by the upregulation of transcriptional drivers including *Runx2* and *Zeb2*, and was followed by later expression of exhaustion markers such as *Havcr2* (Tim-3). As expected *Ctla4* expression was increased in the PAt and Arrested phases (>60◦), consistent with delayed induction. Analysis of genes suppressed by combination therapy (**Figure 2i**) confirmed that the “master regulator” of quiescence, *Bach2* ^39^, was significantly downregulated along the entire trajectory in the treatment group relative to IgG. Similarly, suppression of *Zfp36l1*, an RNA-binding protein known to destabilise cytokine mRNAs ^40^, was consistent with the increased expression of *Ifng* and *Gzmb* transcripts in the treatment group.

Collectively, these data show that, at this early post-treatment time point, combination therapy is associated with coordinated temporal changes in key gene modules, including increased *Zeb2/Runx2*-associated differentiation programmes and reduced *Bach2/Zfp36l1*-associated stemness-linked restraint, alongside expansion of a persistent effector population.

### CanonicalTockySeq Reveals that Combination Immunotherapy Modifies CD4^+^ T cell Temporal Programme

To determine the mechanism by which the combination immunotherapy (anti-PD-L1 + anti-CTLA-4) reinvigorates the helper T cell compartment, we applied CanonicalTockySeq to CD4^+^ T cells in silico-sorted from the B16 melanoma Nr4a3-Tocky dataset (**Figure 2a**). We first defined the high-dimensional differentiation architecture by projecting CD4^+^ T cells into the Canonical Tocky Space (**Figure 3a**). The resulting manifold revealed a continuous trajectory originating from the “New” (Blue) pole and extending through the “Persistent” (BR) state to a terminal “Arrested” (Red) pole. Cross-validation of the Gradient Tocky Time against the experimental sorting labels (**Figure 3b**) demonstrated the algorithm’s power to infer the gaps between the sorted three Tocky fractions and resolve hidden heterogeneity, realigning them to fill the gaps. The “Blue” and “BR” fractions mapped to early and intermediate timepoints, respectively; the “Red” fraction was clearly deconvolved into two distinct subsets: active “late-persistent” cells ( ≈ 50o) and terminally “arrested” cells (> 80o).

**Figure 3.**
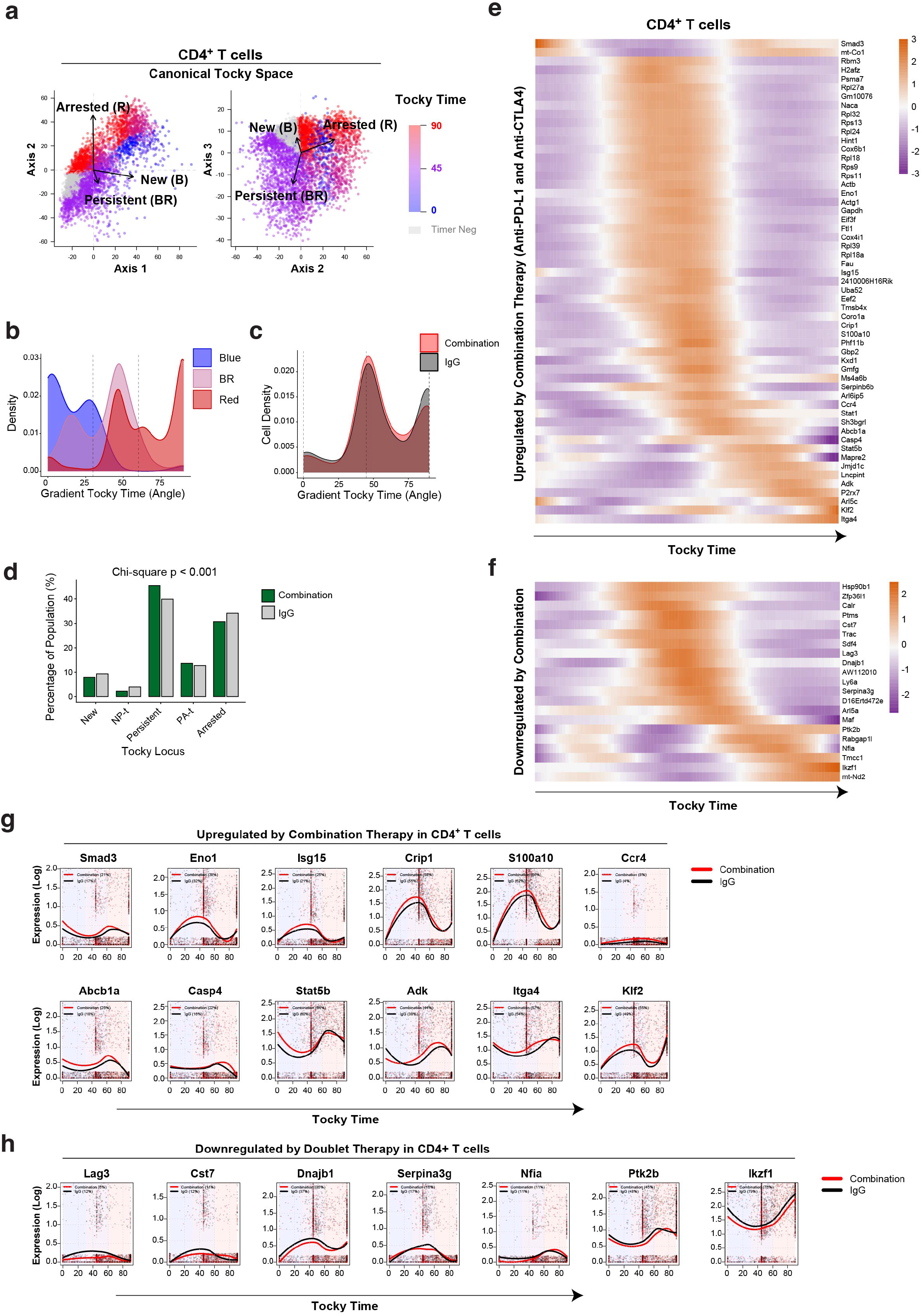
CanonicalTockySeq Analysis of anti-tumour CD4^+^ T cells under checkpoint blockade. **(a)** Multi-Dimensional Canonical Tocky Space Analysis: Projection of CD4^+^ TILs onto the first three canonical axes. Explanatory biplot vectors represent the reference Tocky signatures: New (B), Persistent (BR), and Arrested (R). Single cells are color-coded by their calculated Tocky Time (Angle) on a scale from 0° (New) to 90° (Arrested). (b) Validation of Trajectory vs. Experimental Fractions: Density plots showing the distribution of experimentally sorted Blue, BR, and Red Tocky fractions along the calculated Gradient Tocky Time (Angle). (c) Comparative Cell Density Analysis: Global distribution of CD4+ TIL population density along the Tocky Time trajectory for the combination-treated (red) and IgG control (grey) groups. (d) Population Distribution across Tocky Loci: Bar chart quantifying the percentage of the CD4+ T cell population residing within discrete Tocky Loci: New, NP-t, Persistent, PA-t, and Arrested for both treatment conditions. (e–f) Heatmap Visualization of Transcriptional Cascades: Temporal heatmaps showing the expression of genes significantly upregulated (e) or downregulated (f) by combination therapy. Genes are ordered on the y-axis according to their peak expression timing along the Tocky Time trajectory (x-axis). (g–h) Kinetic Expression Profiles: High-resolution line graphs displaying the temporal expression patterns of representative genes upregulated (g) or downregulated (h) by combination therapy. Individual single cells are plotted along Tocky Time, with red lines representing the treatment group and black lines representing the IgG control.

Quantitative analysis of the Tocky Time density revealed moderate shifts in the Tocky-Time gradient by cell density analysis (**Figure 3c**). This was further supported by the Tocky Locus approach (**Figure 3d**), where a Chi-square test confirmed a non-random redistribution of T cells across temporal phases (*p* < 0.001). Analysis of the CD4^+^ T cell activation cascade (**Figure 3e**) identified a coordinated transition from early signalling to late-stage tissue-homing potential. The programme initiated with the induction of *Smad3*, followed by dominance of translational and glycolytic genes (e.g., *Rps/Rpl* family, *Gapdh*). This metabolic surge is followed by interferon-responsive genes (Stat1, *Isg15*) and culminates in a persistence and homing programme (*Stat5b, Klf2, Itga4*).

Conversely, the silencing cascade (**Figure 3f**) highlighted genes associated with quiescence and restriction, ER-stress regulation, and inhibitory signalling that are actively repressed by the therapy. The silenced programme initiated with the early downregulation of protein-folding chaperones (*Hsp90b1, Calr*), followed by intermediate suppression of the inhibitory checkpoint *Lag3* and the memory marker *Ly6a*. This de-repression phase culminated in silencing of lineage-restricting transcription factors *Maf* and *Ikzf1*.

Gene-by-gene expression plots along the temporal trajectory (**Figure 3g**) provided high-resolution support for these temporal patterns. Glycolytic regulators (e.g., Eno1) and signalling mediators (e.g., *Smad3*) were induced early and maintained at higher levels in the treatment group than in controls. *Stat5b* was markedly upregulated, with a peak in the New and PAt-to-Arrested phases, consistent with rapid induction and sustained representation along the trajectory. *Isg15* and *Crip1* showed pronounced upregulation in the Persistent window, defining this compartment as a distinct activated effector state. In parallel, exhaustion-associated signalling was attenuated, with strong downregulation of Lag3 across the trajectory in combination-treated cells versus IgG controls (**Figure 3h**).

Collectively, these data show that combination therapy preserves global Tocky-Time trajectory structure while extensively remodelling time-resolved gene-regulatory programmes in CD4^+^ T cells.

### Tocky-Time Analysis of Human Melanoma Patient Data Reveals Temporal Programme of Anti-Tumour T Cells Controlled by Checkpoint Blockade

To investigate the temporal dynamics of T cell responses under checkpoint blockade, we projected the scRNA-seq data of CD8^+^ T cells from melanoma patients ^32^ into a canonical Tocky space using the murine Nr4a3-Tocky signature data (**Figure 4a**). We used the scRNA-seq dataset from human melanoma patients under either anti-PD-1 monotherapy or the combination therapy using anti-PD-1 and anti-CTLA-4, which includes clinical response data ^32^. In contrast to the murine experiment, in which samples were analysed 2 days after the final dose within an 18-day treatment schedule (Figures 2 and 3), these human biopsies were obtained at later and heterogeneous timepoints after treatment initiation (first post-treatment biopsy: mean ∼209 days, median 75 days from baseline ^32^) and therefore capture treatment-associated temporal-state structure rather than an immediate post-treatment signalling response.

**Figure 4.**
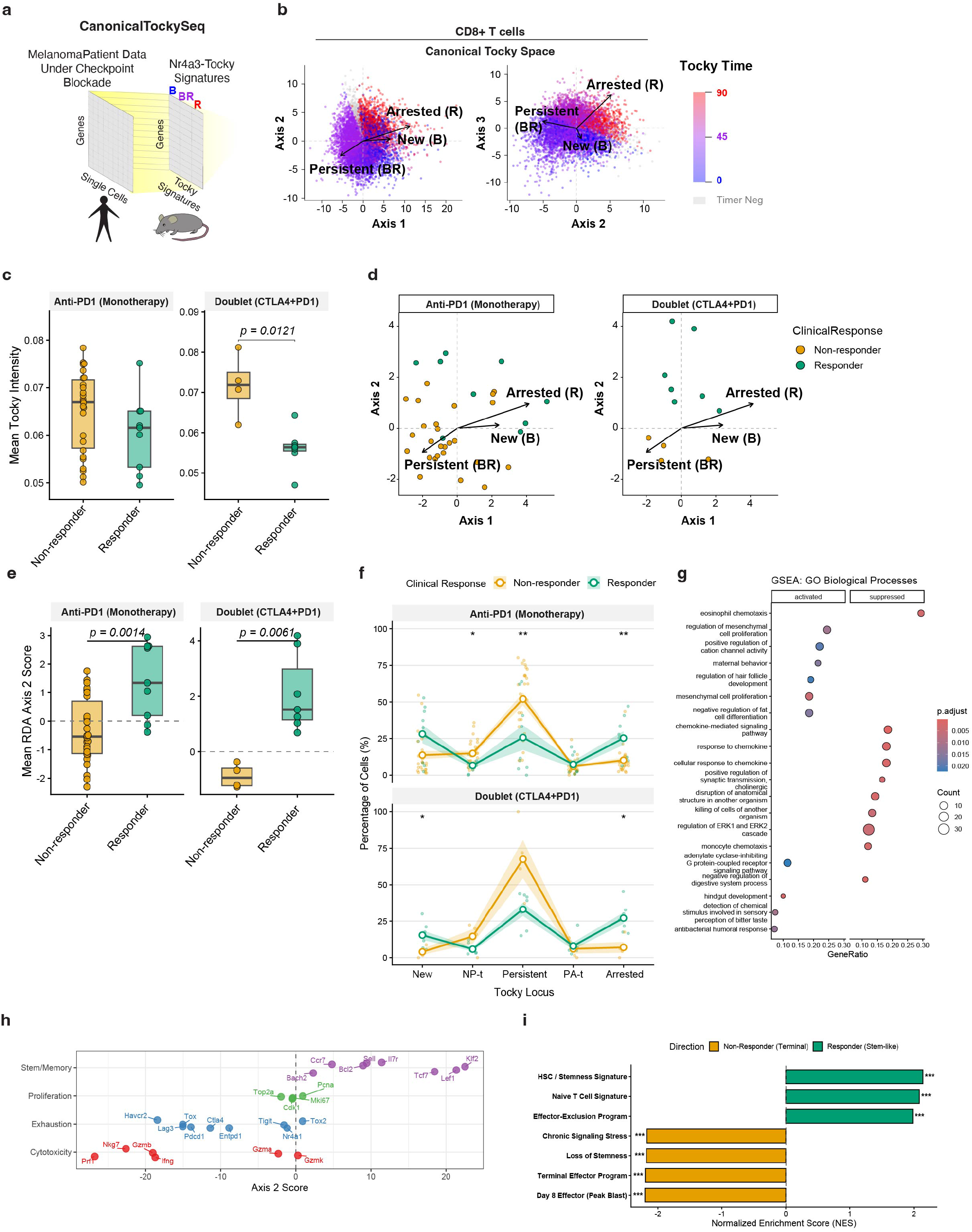
Application of CanonicalTockySeq to Human Melanoma Patient scRNA-seq Data. (a) Schematic of Cross-Species Trajectory Projection. Experimentally defined murine Nr4a3-Tocky signatures (New, Persistent, Arrested) are used to constrain and project human melanoma patient scRNA-seq data into a standardised temporal manifold. (b) Multi-Dimensional Canonical Tocky Space. Projection of human CD8+ T cells onto the first three canonical axes. Explanatory biplot vectors represent the New (B), Persistent (BR), and Arrested (R) signatures. Cells are color-coded by their calculated Tocky Time (Angle), ranging from 0° (New) to 90° (Arrested). (c) Patient-Level Tocky Intensity Analysis. Boxplots comparing the mean Normalised Tocky Intensity between non-responders (orange) and responders (teal) across two treatment cohorts: Anti-PD1 (Monotherapy) and combination (Anti-CTLA4 and anti-PD1). Each dot represents the mean value for an individual patient. (d) Barycentric Patient Distribution. Visualisation of individual patient centroids (mean coordinates) within the 2D Canonical Tocky space (Axis 1 vs. Axis 2), stratified by treatment group and clinical response. (e) Quantitative Analysis of Canonical Tocky space Axis 2. Comparison of mean RDA Axis 2 scores between clinical response groups. Statistics derived from Wilcoxon rank-sum tests performed at the patient level. (f) Tocky Locus Distribution. Quantitative analysis of population flux across the five Tocky stages (New, NP-t, Persistent, PA-t, Arrested). Plots show the percentage of cells within each locus for non-responders (orange) and responders (teal). Statistical significance (* p < 0.05, **p < 0.01) is indicated for specific loci. (g) Functional Pathway Enrichment (GO Biological Processes). Dot plot showing Gene Ontology (GO) terms significantly associated with the Tocky trajectory, categorized by those activated or suppressed along the temporal manifold. (h) Functional Mapping of Gene Signatures on RDA Axis 2. Distribution of canonical genes associated with Stem/Memory, Proliferation, Exhaustion, and Cytotoxicity across the Axis 2 spectrum. Positive scores correlate with responder-associated trajectories, while negative scores correlate with non-responder trajectories. (i) Gene Set Enrichment Analysis (GSEA). Bar chart showing the Normalised Enrichment Score (NES) for specific immunologic and differentiation signatures. Bars represent the enrichment of specific programmes in responders (teal, positive NES) versus non-responders (orange, negative NES). Significance is indicated by asterisks (***p < 0.001).

CanonicalTockySeq successfully identified a Tocky-Time manifold within the human melanoma dataset (**Figure 4b**). This approach allowed us to quantify “Tocky Time” (0–90°), identifying Tocky-equivalent human single cells and the gene profile dynamics downstream of TCR signalling. We next examined TCR signalling intensity within the Tocky manifold model and found that non-responders were characterised by significantly higher mean Tocky intensity compared to responders, particularly in the combination cohort (p = 0.0121, **Figure 4c**). Visualisation of the patient-level centroids within the Canonical Tocky space revealed that responders consistently trended toward Axis 2 positive, which is correlated positively by the Arrested vector, while non-responders clustered toward the Persistent vector (**Figure 4d**). This divergence was captured by the Axis 2 Canonical Tocky score: responders exhibited significantly higher scores in both the anti-PD1 monotherapy and anti-PD1 plus anti-CTLA-4 combination therapy groups (**Figure 4e**).

To unravel these dynamics in the Tocky-Time dimension, we used the Tocky Locus approach and thereby quantitatively captured the features of the responders. T cells from non-responders accumulated heavily in the intermediate Persistent stage, representing active TCR engagement, whereas responders successfully transitioned into the Arrested stage (*p* < 0.01), which indicates the signature of T cells immediately following the cessation of antigen engagement (**Figure 4f**).

Functional mapping of genes driving Canonical Tocky Axis 2 revealed a stark dichotomy between clinical outcomes as well as the associated transcriptional programmes. The responder-associated signature (positive Axis 2) was featured by gene modules consistent with post-stimulation adaptation, including attenuation of ongoing chemokine/ERK-linked signalling and reduced representation of persistent TCR-engagement programmes (**Figure 4g**). The focused analysis of well-characterised genes unravelled that the Canonical Tocky-derived responder-associated signature (positive Axis 2) was defined by “stem-like” and quiescence markers, including *Klf2, Lef1*, and *Tcf7*. In contrast, the non-responder signature (negative Axis 2) was dominated by hyper-cytotoxic effector genes such as *Prf1, Gzmb*, and *Nkg7*, alongside classic exhaustion markers including *Pdcd1, Lag3, Havcr2, CTLA4*, and *Tox* (**Figure 4h**).

This dichotomy was independently supported through Gene Set Enrichment Analysis (GSEA). Responders demonstrated significant enrichment for hematopoietic stem cell (HSC), lymphoid progenitor, naive T cell, and progenitor signatures, confirming the preservation of a progenitor-like renewal transcriptional programme. Conversely, non-responders were enriched for terminal effector signatures, including effector gene sets derived from an activated CD8^+^ T-cell dataset ^41^, as well as signatures associated with stress-responses ^42^ and with stemness loss or lineage priming from an HSC dataset ^43^ (**Figure 4i**).

Taken together, these results suggest that clinical failure is driven by an aberrant diversion of T cells into an excessively-activated, terminally exhausted state that lacks the longevity and self-renewal capacity found in responders. Collectively, our investigation is supportive of the fact that the Canonical Tocky approach with the experimentally-anchored Tocky landmark data unravels temporal mechanisms of anti-tumour response in human T cells and how immune checkpoint blockade can modify them.

## Discussion

The application of the Canonical Tocky approach reveals that T-cell fate is determined not merely by static transcriptional programmes, but by temporally coordinated transcriptional changes that shape differentiation states. In this view, the classically defined “progenitor-like” and “terminally exhausted” states can be further refined by incorporating time-resolved gene regulation linked to TCR signalling dynamics, encompassing both ongoing activity and recent signalling history as measured by Tocky, rather than treating them as fixed lineages. Within this framework, progression along the Tocky-Time continuum reflects the cumulative history of antigen engagement, and ICB actively reprogrammes how T cells interpret and respond to these signalling dynamics over time.

In non-responders, CD8^+^ T cells are enriched in the Persistent Locus, consistent with a differentiation constraint under chronic stimulation. This locus provides a temporal reference for exhaustion-associated commitment ^30, 44^, and places TOX-centred models in a kinetic context ^25, 45^. While TOX is a central regulator of exhaustion, our data suggest that TOX-associated phenotypes are time-resolved states linked to sustained TCR signalling but limited functionality. Mapping TILs to this interval highlights an important dissociation: cells show high downstream TCR-signalling activity (e.g., Tox, Nr4a1, ERK-associated programs) while exhibiting reduced functional progression. This suggests that these markers primarily report a signalled state rather than functional competence. Accordingly, exhaustion in the tumour microenvironment is consistent with a kinetic blockade in which cells remain chronically signalling but transition inefficiently toward resolving trajectories.

This temporal framework also helps reconcile the murine and human datasets. In the murine Nr4a3-Tocky model, analysed 2 days after the final treatment dose, checkpoint blockade was associated with increased representation of CD8^+^ T cells in the Persistent Locus, consistent with sustained antigen engagement and ongoing downstream TCR signalling in the early post-treatment phase. By contrast, the later human melanoma biopsies suggest that, over time, clinically responding T cells are less dominated by persistent antigen engagement, whereas in non-responders CD8^+^ T cells remain enriched in the Persistent Locus, consistent with prolonged antigen signalling and limited transition to later trajectory states. This pattern is compatible with the differentiation constraint described in chronic infection and advanced cancer, where sustained TCR input is coupled to restricted progression toward durable effector or memory-like fates^30, 44^. Together, these temporal patterns suggest that ICB initially increases the representation of antigen-engaged T cells, but that over time a selective process operates within this population, with favourable clinical responses associated with progression away from persistent engagement and toward the Arrested Locus as cells disengage from antigen.

In contrast to cells occupying the Persistent Locus, the progenitor-like compartment occupied earlier temporal trajectory regions relative to antigen exposure. In our dataset, *Bach2* and *Zfp36l1* were enriched in these early regions, whereas, under combination ICB, we observed earlier induction of *Gzmb/Ifng* together with reduced Bach2 expression. These patterns are consistent with a shift from early-state and quiescent programmes ^39, 46^ toward effector-associated programmes ^47, 48^ along the trajectory. Within this framework, Axis 2 summarises this temporal organisation: higher values map to early/quiescence-associated regions, whereas lower values map to Persistent effector and exhaustion-associated regions. This time-resolved organisation is biologically consistent with the established progenitor-to-terminal differentiation hierarchy ^49, 50^.

ICB-associated temporal remodelling is not restricted to CD8^+^ T cells. In CD4^+^ T cells, we observed smaller shifts in trajectory density but clear reorganisation of time-ordered transcriptional programmes. Combination treatment increased expression of survival and metabolic regulators (including *Stat5b* and *Eno1*) along defined trajectory intervals, consistent with altered helper-state kinetics rather than wholesale redistribution of cell states. In parallel, broad reduction of *Lag3* expression across the trajectory suggests delayed or attenuated acquisition of exhaustion-associated transcriptional features. Together, these findings indicate that CD4^+^ responses are modified primarily through temporal rewiring of regulatory cascades within an existing differentiation topology.

CanonicalTockySeq provides an experimentally constrained framework for temporal inference by anchoring transcriptomic structure to biologically defined Tocky landmarks (B/BR/R). Modelling progression with piecewise geodesic interpolation (SLERP) stabilises angular ordering under skewed landmark geometry and decouples temporal position (angle, θ) from signalling magnitude (radial intensity). This decoupling is particularly important in tumour settings, where activation strength and differentiation timing can diverge. Projection of human melanoma data into a murine-anchored temporal framework further supports the cross-context utility of this approach and enables direct testing of how therapies reshape immune dynamics over time, rather than only altering endpoint cell-state composition.

A current limitation of CanonicalTockySeq is the experimental burden of flow-cytometric fractionation and droplet-based single-cell sequencing, particularly because individual Tocky fractions can be rare. Nonetheless, the cross-species transferability shown here argues for the value of establishing standardised Tocky landmark reference datasets. As such resources are expanded and shared, the field will be better positioned to incorporate temporal information into human T-cell analyses and refine interpretation of immune-state dynamics.

## Materials and Methods

### Implementation of Canonical Tocky Analysis

The ordination was performed using a specialised Canonical Redundancy Analysis (RDA) architecture designed for single-cell Tocky data (**Supplementary Methods**). In this gene-centric ordination, genes are treated as the primary observational units (‘sites’) and single cells as the response variables (‘species’), constrained by the explanatory Tocky-defined developmental stages (Blue, Blue-Red, and Red), following the original architecture of RDA ^37^.

This supervised, constrained ordination follows the two-step procedure of multiple linear regression followed by principal component decomposition. The analysis yields two distinct sets of cell coordinates. (1) Fitted Cell Scores represent the idealized cell positions in the “Space of X” (explanatory variables), strictly explained by the Tocky-defined constraints. These scores reflect the rigid, model-driven path of differentiation and tend to be more singular or degenerate due to the imposed constraints. (2) Cell Scores represent the actual cell positions in the “Space of Y” (response variables), derived by projecting the original centred expression data onto the canonical axes.

We utilised the Cell Scores for the final manifold construction to preserve the biological heterogeneity and the natural deviation of individual cells from the idealized model. Biplot vectors were derived by regressing the gene expression scores onto the constraint matrix (z), providing a geometric representation where direction signifies the Tocky time dimension and magnitude represents the collective intensity of the reference signatures (reflecting TCR signaling strength).

### Trajectory Reconstruction via Identification of Tocky Gradient Manifold

To quantify differentiation progress, we developed GradientTockySeq, a deterministic trajectory-projection framework that maps single-cell transcriptomes onto a continuous “Tocky Time” coordinate (**Supplementary Methods**). Rather than assuming linear progression in expression space, GradientTockySeq represents differentiation as directional rotation of transcriptional programmes. The trajectory is defined as a piecewise path connecting three biological landmarks derived from Tocky signatures: New (Blue) → Persistent (Blue-Red) → Arrested (Red). We connected these landmarks using Piecewise Spherical Linear Interpolation (SLERP) ^38^. Because these landmarks vary in Euclidean magnitude, SLERP generates a piecewise logarithmic spiral path across the transcriptomic hypersphere, yielding a smooth curved reference path that accounts for simultaneous shifts in both transcriptional direction and intensity. By tracing rays from the origin through the piecewise SLERP reference path, the model admits a cone-like geometric interpretation.

Single cells were mapped to this manifold by identifying the angle θ (Tocky Time) that maximized the cosine similarity between the cell’s expression vector and the reference trajectory. The derived angle was normalised to a 0–90° scale, where 0° represents the “New” state and 90° represents the “Arrested” state. To distinguish transcriptional volume from differentiation timing, we calculated “Tocky Intensity” as the Euclidean norm (magnitude) of the cell’s vector, normalised against the expected magnitude of the trajectory at angle *θ*. Cells displaying negative correlations to all three landmarks (falling in the quadrant opposite to the Tocky cone) were classified as “Timer Negative” and excluded from trajectory analysis to ensure biological specificity.

### Canonical Tocky Locus Analysis

Gradient Tocky Time (Angle) was partitioned into five 18-degree intervals to define the Tocky Loci: New (0◦ – 18◦), NP-t (18◦ - 36◦), Persistent (36◦ - 54◦), PA-t (54◦ - 72◦), and Arrested (72◦ - 90◦). To evaluate whether the observed distribution of cells across the Tocky-Time loci was independent of the treatment group, a Pearson’s Chi-square test was performed on the raw cell counts.

### Temporal Cascade Analysis and Heatmap Visualisation

To visualise the sequential activation of genes across the Tocky-Time continuum, we used a temporal cascade analysis, which identifies the peak timing of gene expression to reveal the transcriptomic dynamics. The continuous Tocky-Time gradient (0–90°) was discretised into 100 equally sized angular intervals (bins). For each gene, the mean expression was calculated within each bin to generate a raw temporal profile. To reduce stochastic noise and capture the underlying biological signal, these profiles were smoothed using a locally estimated scatterplot smoothing (LOESS) regression (span = 0.5) across the angular grid.

To compare genes with disparate baseline expression levels, the smoothed expression values were Z-score normalized (mean = 0, standard deviation = 1). To ensure visual clarity and minimize the impact of extreme outliers, the scaled values were capped at a range of [-3, 3]. To construct the temporal cascade, for each gene, the Tocky-Time bin corresponding to the maximum Z-score value was identified as the Peak Time, i.e. Peak Time_*g*_ = argmax_*T*_(*z*_*g,T*_). Genes were subsequently ranked and ordered from earliest to latest peak timing. This serial arrangement visualizes the transcriptional dynamics, allowing for the identification of early-, intermediate-, and late-response genes along the temporal gradient.

The resulting cascade was visualised as a heatmap using a divergent colour palette (Purple-White-Orange) using the R package pheatmap ^51^. The x-axis represents the Tocky-Time (0– 90°), and the y-axis represents individual genes sorted by their peak activation.

### Integrated Differential Dynamics and Magnitude Analysis on the Gradient Tocky-Time

To identify treatment-associated transcriptional programmes along T-cell differentiation, we developed a two-stage computational pipeline coupling Gradient Tocky-Time temporal coordinates with Generalized Additive Model (GAM) modelling.

To prioritise genes with the highest information content along the Tocky trajectory, we first performed an initial feature screening using the SelectTockyGenes function. We utilized Gradient Tocky Time (the Tocky Angle) as the independent variable. Genes were filtered based on a minimum expression threshold (detected in >5% of cells), and their temporal dynamics were scored using a rapid linear regression model with natural splines (*df* = 3): *Expression* ∼ *ns*(Tocky Time, *df* = 3). The top 2,000 genes, ranked by their coefficient of determination (*R*^2^), were selected for downstream rigorous GAM fitting.

To identify treatment-associated transcriptional programmes along T-cell differentiation, we coupled CanonicalTockySeq temporal coordinates with a Generalized Additive Model (GAM) modelling using the R package tradeSeq ^52^. Using the output of GradientTockySeq, i.e. Gradient Tocky Time as the pseudotime covariate and we tested whether gene-expression dynamics diverged between experimental conditions, excluding cells with undefined angle values (Timer-negative) from the trajectory model. For each gene, expression was modelled using a negative-binomial GAM with cubic regression splines (nknots = 5) and condition as a covariate. Because the Tocky trajectory defines a single, non-branching temporal axis, we fitted a single lineage with unit weights (*w* = 1) for all constituent cells. To identify treatment-specific effects, we utilized the conditionTest function and p values were adjusted using Benjamini–Hochberg’s false discovery rate. To separate temporal-shape effects from overall abundance effects, we applied a second, directional filtering layer with adjusted p-values < 0.05 and average log2 fold change > 0.1 and ranked them by descending the fold change.

### Data Preprocessing, In silico Sorting, and CanonicalTockySeq Analysis of Human Data

Human melanoma scRNA-seq data (GSE120575 ^32^) was integrated by identifying human orthologs for the murine Tocky signatures. Single-cell TPM (Transcripts Per Million) expression data and clinical metadata from the Sade-Feldman melanoma cohort (GSE120575 ^32^) were retrieved from the GEO database. Data were processed using Seurat (version 4 ^53^). Initial filtering retained cells expressing at least 200 genes and genes detected in at least 3 cells. Following normalisation and scaling, the top 2,000 highly variable features were identified. Dimensionality reduction was performed via Principal Component Analysis (PCA) using the first 10 principal components. To isolate specific T-cell populations, we employed a hierarchical in silico sorting strategy based on canonical marker expression and graph-based clustering (resolution = 1.2). First, clusters were screened for T-cell markers (CD3G, CD3D, CD3E) and filtered to exclude non-T cell lineages (e.g., CD19^+^ B cells or ITGAM^+^ myeloid cells). The remaining CD3^+^ cells were re-embedded and re-clustered. CD8^+^ T cells were defined by clusters exhibiting high CD8A/B and cytotoxic marker expression. CD4^+^ T cells were identified by CD4 expression and included both FOXP3^-^ and FOXP3^+^ T-cell clusters.

To apply the CanonicalTockySeq framework to the human melanoma single-cell dataset using the murine Nr4a3-Tocky data as explanatory variables, we performed a cross-species gene correspondence between Human (HGNC) and Mouse (MGI) symbols using the biomaRt R package ^54^. We queried the Human Ensembl database (`hsapiens_gene_ensembl`) to retrieve the `mmusculus_homolog_associated_gene_name` attribute for all genes detected in the human T-cell subsets. To maintain data integrity, only genes with confirmed orthologous pairs were retained. To prevent signal loss and preserve total transcriptional output, we implemented a summation strategy for many-to-one relationships. After subsetting the human expression matrix to include only mappable genes, we applied a rowsum operation to collapse the expression values of multiple human paralogs into a single “mouse-equivalent” transcriptional unit. We identified the intersection of genes present in both the collapsed human expression matrix and the Tocky signature matrix (n = 15,332). Both matrices were subset and reordered to ensure exact row-wise correspondence. The mouse-converted human matrix data was thus analysed canonically with the Nr4a3-Tocky explanatory data using the CanonicalTockySeq algorithm.

To analyse the output of CanonicalTockySeq, Patient-level aggregation was performed by calculating the mean Tocky Time and Intensity per individual to facilitate statistically robust clinical comparisons. Wilcox test was used for p-values.

### Simulation

Simulation Design We modelled a differentiation path moving through three orthogonal landmark vectors—Blue (*B*, Axis 1), Blue-Red (*BR*, Axis 2), and Red (*R*, Axis 3)—creating a 3D “corner” trajectory that cannot be accurately captured by a single 2D plane. To replicate biological heterogeneity, we introduced two types of variance: (1) Random dispersion (*SD* = 0.2) was added to cell coordinates to simulate transcriptional noise as Gaussian noise; (2) cells were assigned random magnitudes ranging from 0.5 to 4.0. This created a “conical” distribution where cells fanned out from the origin, simulating the wide range of total gene expression intensities (library sizes) observed in real scRNA-seq datasets.

### Mice, Tumour inoculation, and in vivo antibody administration

*Nr4a3*-Tocky mice, or BAC Tg (*Nr4a3*^*ΔExon3 FTfast*^), were generated by the Ono group and reported previously ^6^. Briefly, *Nr4a3*-Tocky was generated by a bacterial artificial chromosome (BAC) approach, in which the Fluorescent Timer (FTfast^55^) gene replaced the first coding exon of the *Nr4a3* gene by a knock-in knock-out approach. Pronuclear injection was used to generate *Nr4a3-Tocky*. The mice were subsequently bred with *Foxp3*-IRES-GFP (C.Cg-Foxp3tm2Tch/J, Jax ID 006769) and maintained in a C57BL/6 background. All animal procedures were conducted in compliance with the UK Animals (Scientific Procedures) Act 1986, under the oversight of the Animal Welfare and Ethical Review Body (AWERB) at Imperial College London, the Animal Ethics Committee at the Institute of Cancer Research, and the Animal Experiment Committee at Kumamoto University.

The B16-F1 melanoma cell line was maintained in DMEM supplemented with 10% FBS, 2 mM L-Glutamine, and Penicillin/Streptomycin. For tumour induction, 2 × 10^5^ cells were injected subcutaneously into the right flank of Nr4a3-Tocky or wild-type C57BL/6 control mice. Immunotherapy was administered via intraperitoneal injection: 0.15 mg of anti-CTLA-4 mAb (clone 9H10, BioXCell) or Syrian Hamster IgG control was administered on days 12, 14, and 16; 0.2 mg of anti-PD-L1 mAb (clone 1-111A) or Rat IgG2A control (clone 2A3, BioXCell) on days 14 and 16. Mice were culled for analysis on day 18 post-inoculation.

### Droplet scRNA-seq Library Preparation and Data Preprocessing

Upon harvesting, tumours were mechanically minced and enzymatically digested in PBS supplemented with Liberase (25 µg/mL), DNase I (250 µg/mL), and 1X Trypsin-EDTA (Sigma-Aldrich) for 10 minutes at 37°C followed by 20 minutes at room temperature under continuous agitation. The resulting tumour suspensions were filtered through a 70 µm mesh and washed with FACS buffer containing 5 mM EDTA. Subsequently, haematopoietic cells were selected using a magnetic sorting method using the mouse CD45 microbeads, LS columns (Milteny Biotech). Simultaneously, tumour-draining lymph nodes (tdLNs) were mechanically dissociated through a 70 µm cell strainer. To enable sample multiplexing and minimise batch effects, tdLN cells and enriched CD45^+^ TILs, each representing pooled samples from multiple mice per biological group, were labelled using the 3’ CellPlex Kit (10x Genomics). Samples were tagged with unique lipid-conjugated oligos as follows: CMO301 (LN-IgG), CMO302 (LN_Dbl), CMO303 (TU_IgG), and CMO304 (TU_Dbl). Following labeling, the multiplexed populations were washed and proceeded to fluorescence-activated cell sorting (FACS) for the isolation of specific Tocky fractions.

To establish the transcriptomic Tocky reference scRNA-seq data required for CanonicalTockySeq, Tocky subpopulations were isolated from the pre-sorted lymphocytes via fluorescence-activated cell sorting (FACS) using a FACSAria (BD Biosciences) into the four Tocky fractions: New (Blue^+^Red^−^), Persistent (Blue^+^Red^+^), Arrested (Blue^−^Red^+^), and Timer negative T-cells. Sorted cells were processed immediately for library construction using the Chromium Single Cell 3’ Library & Gel Bead Kit (v3.1, 10x Genomics). Following quantification via TapeStation (Agilent), high-throughput sequencing was performed on two lanes of a HiSeq X Ten system (Illumina) at the Macrogen Japan NGS facility.

Fastq data were demultiplexed and aligned to the mouse genome (mm10) using the Cell Ranger multi pipeline (10x Genomics) to resolve CellPlex (CMO) assignments. The data were further subject to in silico sorting to purify CD4^+^ and CD8^+^ T cells using the R package Seurat. The CanonicalTockySeq pipeline was applied to the entire pooled transcriptome data using the explanatory vectors derived from the three Tocky landmark populations and thereby to project cells into a three-dimensional Canonical space.

## Acknowledgements

MO was supported by a CRUK Programme Foundation Award (DCRPGF\100007). This research was also supported by KAKENHI research grants from the Japan Society for the Promotion of Science (JSPS) (JP21K07082, JP21H00433, and JP24K10259 to MO; JP25K02693 and JP25K22556 to YS), Japan Agency for Medical Research and Development (AMED) (25gm1810001s0104 to MO; 25gm1810001h0004 and JP223fa627001 to TO; JP23wm0325068 and JP25fk0410070 to YS), and JST-ASPIRE Program (JP29jf0126018). AM was supported by a CRUK Programme Award (CRM183X) and the Institute of Cancer Research/Royal Marsden Hospital Centre for Translational Immunotherapy. KH was supported by CRUK Programme Award A23275, CRUK Programme Award (CRM183X), the ICR/RM CRUK RadNet Centre of Excellence. JH was supported by an ICR studentship from CRUK Convergence Science Centre.

## Author Contributions Statement

MO conceived the study and all computational and Canonical Tocky, and manifold methodologies. MO implemented these as R package modules, authored the analysis code, and conducted all formal analysis and visualisation. JH, MP, SF, LA, TO, YS, KH, AM, and MO designed experiments and developed experimental protocols. MP, SF, and LA provided technical supervision and optimised the B16 in vivo models and experimental protocols. JH, OR, NI, IO, and MO performed experiments. JH, TO, YS, KH, AM, and MO designed experiments. Funding was secured by TO, KH, AM, YS, and MO. JH drafted sections of the biological methods. MO drafted and wrote the entire manuscript.

## Data Availability

The sequencing data reported in this study have been deposited in the NCBI Sequence Read Archive (SRA) under BioProject accession PRJNA1426150.

## Code Availability

The code used to perform CanonicalTockySeq and SLERP manifold construction in this study is publicly available at GitHub:

https://github.com/MonoTockyLab/CanonicalTockySeq

## Supplementary Information

### Supplementary Figure Legends

**Supplementary Figure 1.**
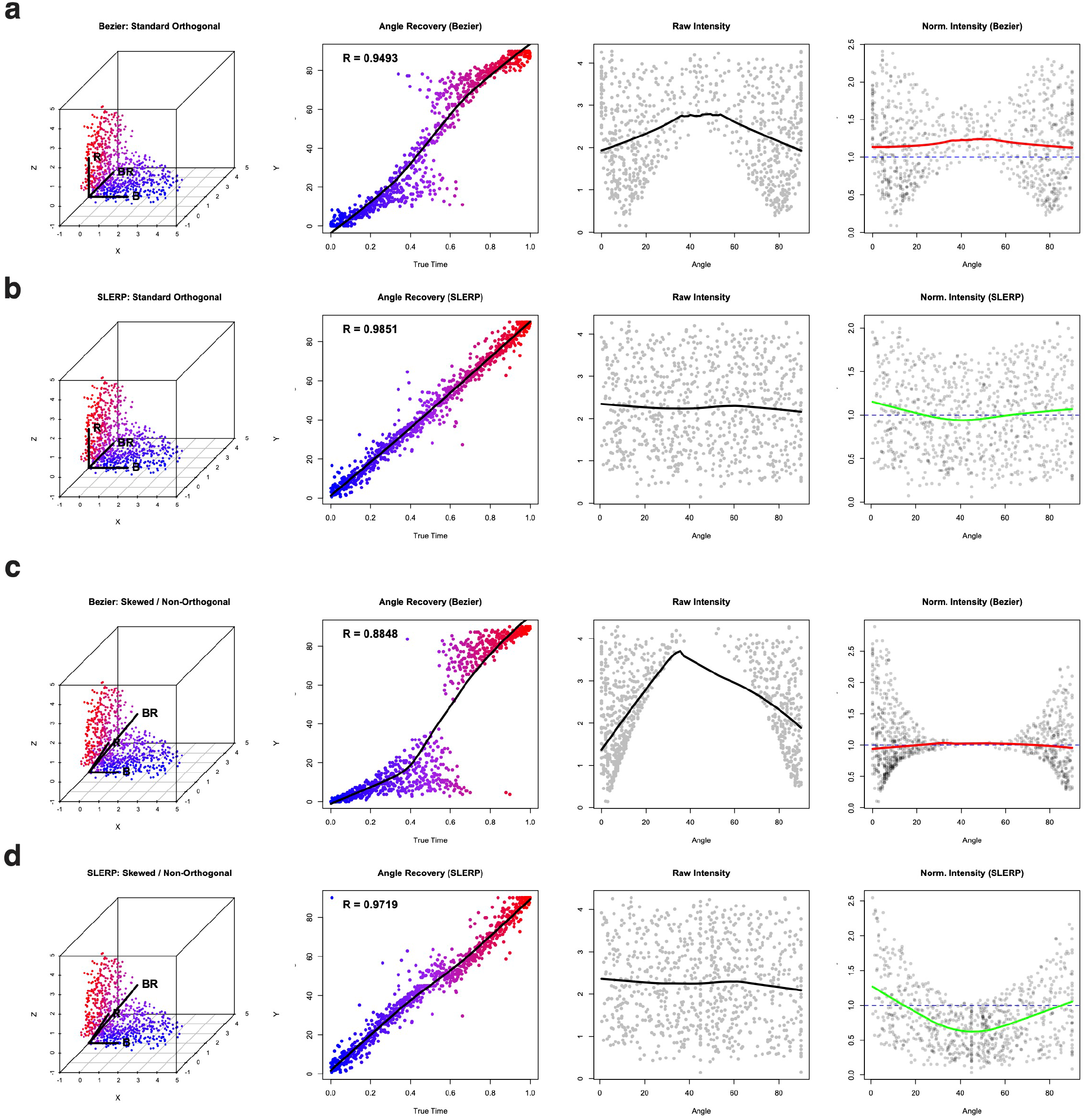
Benchmarking Geometric Validation of GradientTockySeq via 3D Radial Simulation. **a**. Simulation using in orthogonal unit Tocky vectors (B, BR, and R) and manifold identification using Bézier Model. The Angle Recovery plot (column 2) shows slight non-linear deviations from the true simulation time (Pearson r = 0.9493). Raw Intensity (column 3) reflects the simulated radial peak, while Normalized Intensity (column 4) shows the model’s baseline calculation. **b**. Simulation using in orthogonal unit Tocky vectors (B, BR, and R) and manifold identification using SLERP Model. SLERP achieves near-perfect linear angle recovery (r = 0.9851). Raw Intensity (column 3) and Normalized Intensity (column 4) show non-skewed and robust estimation. **c**. Simulation using in skewed Tocky vectors and manifold identification using Bézier Model. This “stress-test” using non-orthogonal landmarks and unequal magnitudes to mimic biological variance. The model fails to maintain geometric fidelity, resulting in a severe sigmoidal “S-curve” artifact in Angle Recovery (r = 0.8848). Raw and Normalized Intensity profiles show significant artifacts (column 4). **d**. Simulation using in skewed Tocky vectors and manifold identification using SLERP Model. Despite the geometric distortion of the ordination space, SLERP maintains robust linear angle recovery (r = 0.9719). Importantly, the Normalized Intensity profile (column 4) remains stable, producing an expected curve given the skewed BR vector longer than the other two vectors.

**Supplementary Figure 2.**
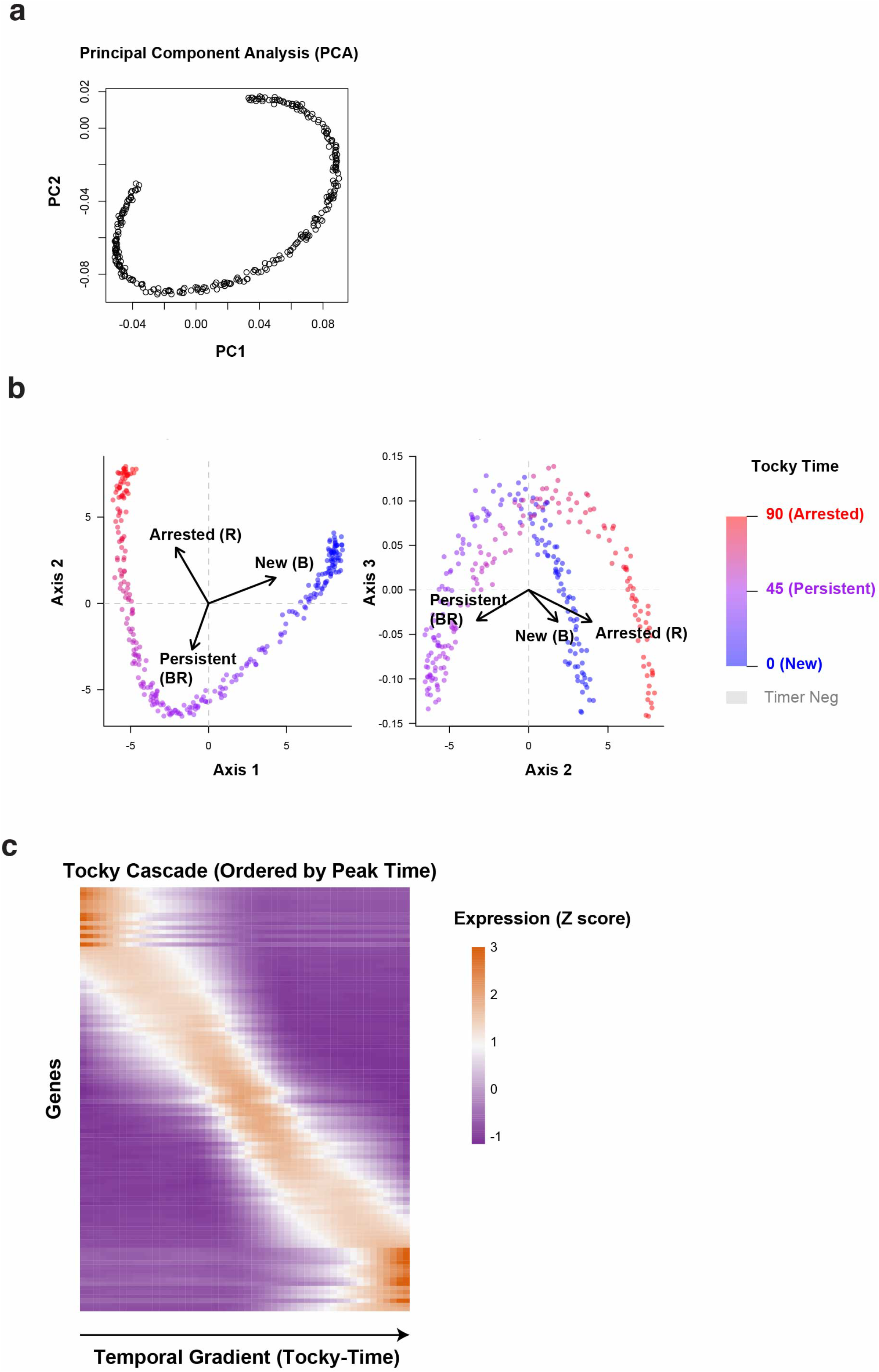
Comparison of Unsupervised PCA and Supervised CanonicalTockySeq Using Simulated Data. A transcriptomic dataset of 300 cells and 500 genes were simulated tomirror biological differentiation, assigning each cell a “true” developmental time (t) ranging from 0 to π/2 . We engineered a continuous transcriptional program by assigning Gaussian tuning curves to the top 100 genes, ensuring that their expression peaks occurred sequentially along the trajectory. Background transcriptional noise was simulated for the remaining 400 genes. The supervised RDA was anchored to three landmark signatures: New (peaking at t = 0), Persistent (peaking at t = π/4), and Arrested (peaking at t = π /2). As shown in the “Getting Started” documentation, the resulting ordination successfully constructed a transcriptomic manifold with distinct conical geometry. The Axis 1 vs. Axis 2 projection revealed a clear, continuous arched trajectory that accurately mirrored the engineered temporal progression from the “Blue” (New) landmark through the “Blue-Red” (Persistent) landmark, concluding at the “Red” (Arrested) landmark. (a) Unsupervised Principal Component Analysis (PCA). PCA resolves the dominant arched manifold structure driven by sequential gene expression peaks but lacks intrinsic biological orientation or temporal coordinates. (b) Supervised CanonicalTockySeq Ordination. The Canonical Redundancy Analysis (RDA) architecture anchors the transcriptomic manifold to biologically defined landmarks: New (B), Persistent (BR), and Arrested (R). This supervision aligns the arched manifold with the developmental progression, allowing for the calculation of continuous Tocky-Time (0–90°). (c) Sequential Transcriptional Cascade Recovery. Gradient analysis using piecewise SLERP accurately resolves the sequential activation of 100 engineered Gaussian tuning curves. The smooth “waterfall” pattern confirms the high temporal resolution of the geodesic interpolation and its ability to faithfully reconstruct developmental sequences.

### Supplementary Methods: Mathematical Framework of CanonicalTockySeq and GradientTockySeq

#### 1. Introduction and Rationale

Experimentally analysing temporal transcriptional dynamics at single-cell resolution in vivo remains a major challenge. Standard single-cell RNA sequencing (scRNA-seq) provides a high-dimensional “snap-shot” of cellular heterogeneity but fails to capture the critical temporal history of T-cell receptor (TCR) engagement. While computational frameworks such as pseudotime trajectory analysis and RNA velocity attempt to infer cellular progress, they are inherently limited by the lack of an absolute temporal anchor and can be sensitive to assumptions that are challenged by rapid transcriptional and metabolic remodelling during T-cell activation.

To address these constraints, we developed **CanonicalTockySeq**, a framework that integrates a molecular clock of TCR signalling, the Nr4a3-Tocky system [1], with scRNA-seq to establish an experimentally anchored temporal reference. The Nr4a3-Tocky system utilises a Fluorescent Timer protein that matures from an unstable Blue-fluorescent form (*t*_1*/*2_ ≈ 4 h) to a stable Red-fluorescent form (*t*_1*/*2_ ≈ 122 h) [2]. This biophysical process allows for the stratification of cells into distinct developmental stages: *New* (Blue), *Persistent* (Blue-Red), and *Arrested* (Red).

In this framework, we use flow-sorted Tocky populations as experimentally defined biological landmarks. By treating these distinct populations as landmarks, CanonicalTockySeq constructs a transcrip-tomic manifold in canonical space with conical geometry. This approach effectively separates temporal progression (encoded as the geodesic angle, or Tocky-Time) from signalling strength (encoded as radial intensity), enabling the time-resolved analysis of gene-expression dynamics in vivo.

This Supplementary Methods section details the mathematical and algorithmic formulation of the two core components of this framework: `CanonicalTockySeq` (dimensionality reduction and canonical space definition) and `GradientTockySeq` (manifold construction and trajectory quantification).

##### 1 CanonicalTockySeq

The `CanonicalTockySeq` function implements Redundancy Analysis (RDA), a supervised canonical ordination method, to order single cells along a differentiation trajectory explicitly constrained by Tocky timer fluorescence.

###### 1.1 Biological Constraints and Data Structure

Unlike unsupervised dimensionality reduction methods (e.g., PCA or UMAP), CanonicalTockySeq is a supervised framework. We used the sorted Nr4a3-Tocky landmark populations, **New** (*B*), **Persistent** (*BR*), and **Arrested** (*R*), as explanatory variables.

Let the input data be defined as:

- **X** ∈ ℝ^*G×C*^ : The Gene Expression Matrix, where *G* is the number of genes (rows) and *C* is the number of cells (columns).
- **Z** ∈ ℝ^*G×K*^ : The Explanatory (Constraint) Matrix, where *K* is the number of Tocky signatures (here *K* = 3 corresponding to the *B, BR*, and *R* stages).

###### 1.1.1 Architectural Interpretation: Gene-Centred Ordination

In standard multivariate ecology (e.g., `vegan::rda` [3]), rows typically represent independent samples (Sites). In `CanonicalTockySeq`, we invert this architecture to create a gene-centric ordination:

- **Rows (Genes):** Treated as the fixed features defining the constrained structure.
- **Columns (Cells):** Treated as the entities projected into this structure.
- **Constraints (Z):** Describe gene-level association with Tocky stages derived from the sorted landmark populations.

###### 1.2 Mathematical Algorithm

The function performs RDA via Singular Value Decomposition (SVD) on a matrix of fitted values obtained through multivariate linear regression.

###### 1. Standardisation

To emphasise relative transcriptional profile shape rather than absolute expression magnitude, we preprocess the expression matrix **X** and constraint matrix **Z** column-wise. Because **X** is organised as genes *×* cells, column-wise standardisation centres and scales each individual cell profile across genes:

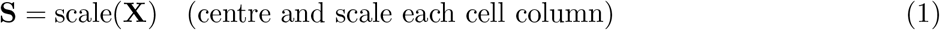

The Tocky constraint matrix is scaled column-wise without centring:

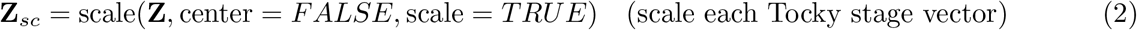

This transformation reduces differences in overall expression magnitude between cells and focuses the constrained ordination on relative transcriptional pattern, while preserving the directional structure of the Tocky stage vectors.

###### 2. Constrained Ordination (Projection)

We seek to explain the variance in cell profiles (**S**) using linear combinations of the Tocky gene signatures (**Z**_*sc*_). We compute the projection matrix (hat matrix) using ordinary least squares (OLS):

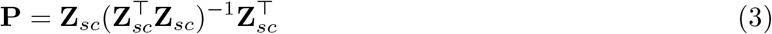

The matrix of fitted values (**S**^∗^) represents the component of the standardised expression matrix captured by the Tocky-defined constraint space and is calculated as:

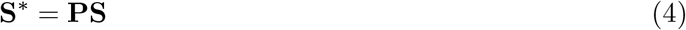

###### 3. Dimensionality Reduction (Partial SVD)

We perform SVD on the constrained matrix **S**^∗^ to identify the principal axes of variation within the Tocky-defined subspace. Using the `irlba` algorithm for computational efficiency, we compute the top *K* components:

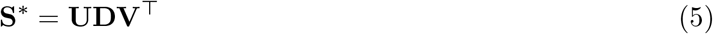

Where:

- **U** ∈ ℝ^*G×K*^ : Left singular vectors (Gene Loadings).
- **D** ∈ ℝ^*K×K*^ : Diagonal matrix of singular values.
- **V** ∈ ℝ^*C×K*^ : Right singular vectors (Cell Loadings).

###### 1.3 Score Calculation and Biological Interpretation

The algorithm outputs coordinates that allow for the simultaneous visualisation of cells, genes, and Tocky stage constraints.

###### 1. Cell Scores

These represent the actual positions of the cells in the ordination space. They are calculated by projecting the original standardised data **S** onto the gene axes **U**. This captures biological heterogeneity not fully explained by the Tocky constraints:

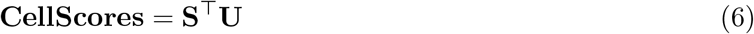

Cell scores are subsequently centred to ensure the trajectory is relative to the population mean.

###### 2. Fitted Cell Scores

These represent the idealised positions of cells under the linear transitions defined by the Tocky vectors:

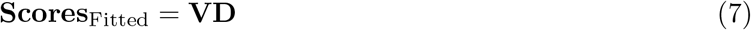

###### 3. Gene Scores (Expression Scores)

Gene scores quantify the loading of each gene on the canonical axes, representing the contribution of each gene to the Tocky-defined manifold. In ordination space, these scores provide the coordinates of gene anchors relative to the cellular trajectory:

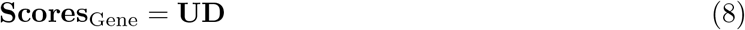

###### 4. Biplot Scores

Biplot scores map the original Tocky constraints (**Z**) directionally into the canonical space. Unlike the constrained ordination step, which uses the scaled constraint matrix **Z**_*sc*_, biplot coordinates are obtained by regressing the gene scores on the original constraint matrix **Z** via ordinary least squares:

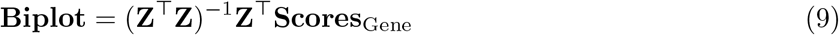

The resulting vectors indicate the association of each Tocky stage signature with the canonical axes.

#### 2 GradientTockySeq

While CanonicalTockySeq constructs the manifold, quantification requires a continuous metric. `GradientTockySeq` constructs a deterministic Spherical Linear Interpolation (SLERP) Manifold within this constrained space to estimate the continuous temporal transition of cells.

To convert manifold geometry into quantitative temporal metrics, we modelled the progression as a directional rotation of transcriptional programmes. We utilised Piecewise SLERP [5] to generate a smooth curved path between the Tocky landmarks. The manifold is anchored by raw landmark vectors corresponding to the **New** (*B*), **Persistent** (*BR*), and **Arrested** (*R*) signatures.

Let the raw landmark vectors be defined as:

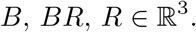

Unlike standard spherical projection, `GradientTockySeq` preserves the relative magnitudes of these landmarks, generating a spiral-like trajectory on a conical manifold rather than a fixed-radius hypersphere. This allows for the decoupling of **Tocky Time** (angular progression) from **Tocky Intensity** (radial magnitude).

##### 1. Timer-negative pre-filtering

For each cell *x*_*i*_ ∈ ℝ^3^, projections onto the landmark axes were computed:

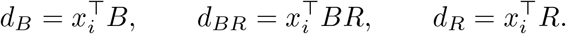

Cells falling into the quadrant opposite the differentiation cone were identified. If `filter negative = TRUE`, cells satisfying

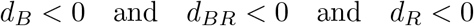

were classified as *Timer-negative* and excluded from trajectory mapping to ensure biological specificity.

##### 2. Piecewise SLERP-weighted trajectory construction

First, the angular distances between landmark *directions* were computed using unit-normalised vectors:

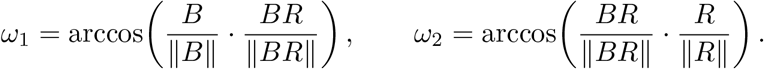

Next, a continuous piecewise trajectory was generated. SLERP-derived angular interpolation weights were applied to the *raw* landmark vectors *B, BR*, and *R*, allowing the reference path to reflect both directional change and landmark magnitude. This produces a curved trajectory on a conical manifold rather than a fixed-radius hypersphere. For a local parameter *h* ∈ [0, 1]:

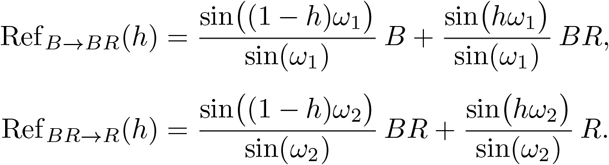

##### 3. Cell-to-trajectory mapping (Tocky Time)

To map cells onto this conical manifold, we solved for the optimal trajectory point that maximises angular similarity. For each valid cell *x*_*i*_, we optimised *h* ∈ [0, 1] to maximise the cosine similarity (equivalent to maximising the normalised projection):

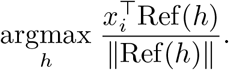

This formulation projects cells onto the “rays” defined by the trajectory, effectively decoupling angular progression from radial intensity. The optimal local *h* was converted to a global coordinate *t*_*i*_ ∈ [0, 1]:

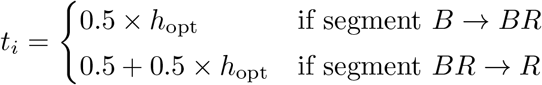

The final **Gradient Tocky Time** (Angle) is reported as:

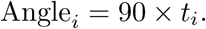

##### 4. Intensity quantification

Signal intensity was quantified via the Euclidean norm of the cell vector:

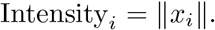

To normalise for the varying magnitude of the trajectory (the conical shape), we calculated the expected magnitude at the assigned time *t*_*i*_:

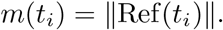

The **Normalised Intensity** represents the radial deviation from the manifold surface, reflecting signature strength relative to the expected baseline at that specific temporal stage:

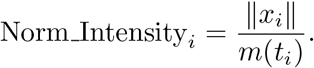

